# Electrophysiological Representations of Multivariate Human Emotion Experience

**DOI:** 10.1101/2023.05.23.541871

**Authors:** Jin Liu, Xin Hu, Xinke Shen, Sen Song, Dan Zhang

**Author notes:** D. Zhang is the corresponding author. D. Zhang is with the Department of Psychology and with the Tsinghua Laboratory of Brain and Intelligence, Tsinghua University, Beijing, China, 100084.

## Abstract

Despite the fact that human daily emotions are co-occurring by nature, most neuroscience studies have primarily adopted a univariate approach to identify the neural representation of emotion (emotion experience within a single emotion category) without adequate consideration to the co-occurrence of different emotions (emotion experience across different emotion categories simultaneously). To investigate the neural representations of multivariate emotion experience, this study employed the inter-situation representational similarity analysis (RSA) method. Researchers used an EEG dataset of 78 participants who watched 28 video clips and rated their experience on eight emotion categories. The EEG-based electrophysiological representation was extracted as the power spectral density (PSD) feature per channel in the five frequency bands. The inter-situation RSA method revealed significant correlations between the multivariate emotion experience ratings and PSD features in the Alpha and Beta bands, primarily over the frontal and parietal-occipital brain regions. The study found the identified EEG representations to be reliable with sufficient situations and participants. Moreover, through a series of ablation analyses, the inter-situation RSA further demonstrated the stability and specificity of the EEG representations for multivariate emotion experience. These findings highlight the importance of adopting a multivariate perspective for a comprehensive understanding of neural representation of human emotion experience.

## Introduction

Human emotion experience is complex and hybrid, often involving the co-occurrence of multiple emotions (Ellsworth et al., 1988). For instance, people may experience a mixture of sadness and joy during their graduation ceremony, but feel fear and disgust when they see a mouse in the room. Researches conducted in real-life situations (e.g., sport competition) have shown that people often experience two or more emotion categories simultaneously (Martinent et al., 2012). While the resulting emotion experience could be described as the combination of multiple co-occurring emotion categories (or simply termed as the ‘mixed emotions’, which was defined as the co-occurrence of any two or more same-valence or opposite-valence emotions (Scherer, 1998)), it could also be considered as a multivariate^1^ construct with each emotion category as one variate. The ‘multivariate’ as a terminology of mathematical modelling is used to characterize the instance with complex structure specifically, so the multivariate construct could be advantageous in describing fine-grained emotion experience. For instance, different emotional situations such as the graduation ceremony and wedding ceremony, might be associated with different levels of mixture of sadness and joy. In line with the empirical observations, it has been reported that the self-report emotion experience with rich emotion categories could be an effective and reliable tool for the assessment of emotional reactivity and regulation, as reflected by the Multidimensional Emotion Questionnaire (MEQ) (Klonsky et al., 2019). Meanwhile, the co-occurrence patterns of different emotions have been suggested to be related to appraisals and action tendencies (Miyamoto et al., 2010) as well as psychological well-beings (Adler et al., 2012).

In the past two decades, the experimental psychology researches on mixed emotions have been emerging, continuing to shed light on the structure of affect, and the state-of-the-art mixed emotion studies mainly focused on the co-occurrence of positive and negative affects (such as happiness and sadness (Larsen et al., 2014)), with greater understanding of developmental, cultural, and individual differences. For example, it has been reported that older children were more likely than younger children to actually experience both happiness and sadness when watching bittersweet films (Larsen et al., 2007), indicating the developmental changes in the understanding, and experience of emotional contradiction. However, there are some limitations and further researches are needed. On one hand, the co-occurrence of same-valence emotions also deserves attention and should not be ignored. In terms of correlations, emotions with the same valence tended to be positively correlated (such as the co-occurrence of fear and disgust was common in daily life). In addition, it has been found that the diversity of positive / negative emotions plays an important role in both mental health and physical health, highlighting the benefits of having a rich and complex emotional life (Quoidbach et al., 2014). On the other hand, the emotion categories included in these studies were limited (usually no more than two), leading to the difficulty to portray the multivariate emotion experience in detail. For a more balanced and comprehensive understanding of the nature of mixed emotions, in this study, we intended to include more complex and delicate emotional situations (e.g., the stimuli could evoke multivariate emotion experience as rich as possible), to explore the physiological substrates behind the mixed emotions.

Despite the wealth of behavioral studies on mixed emotions, little is currently known about the neural basis of the co-occurring human emotions. Previous studies on the neural representations of emotion have often overlooked the co-occurring nature of human emotions. Some studies may assume that participants could exclusively feel the certain discrete emotion as intended by the researchers (Hamann et al., 2012), despite the fact that even standard stimuli designed for eliciting discrete emotions could not guarantee the complete absence of other non-target emotions (Hu et al., 2022). Others might presume that the coexistence of non-target emotions would not interfere with the neural representations of the target emotions. Nevertheless, such assumptions may not always hold. For example, it has been found that anger evoked in a social context was usually blended with anxiety and associated with the distinct frontal cortical activity from traditional anger researches, which reflected its withdrawal (vs. approach) motivational tendency (Zinner et al., 2008). Thus, it is suggested that adopting a holistic perspective of multivariate emotion experience may provide new insights into the neural mechanisms underlying emotions. Although this issue is currently under-investigated, it is expected that exploring the multivariate structure of emotions will provide a more complete picture of our emotion experience compared to analyzing single emotion individually (the univariate emotion experience) (Larsen et al., 2001). A recent study analyzed cardiorespiratory physiological recordings in daily life and found that individuals with higher emotional granularity showed a larger number of distinct patterns of physiological activity during resting conditions and more situation-specific activity during emotional events (Hoemann et al., 2021). Another recent EEG study utilized an inter-subject representational similarity analysis method to make full use of the emotion experience with multivariate structure, rather than reducing it to a univariate form. This study identified the involvement of both the frontal and temporo-parietal regions in representing individual differences in human emotions (Hu et al., 2022). Although these studies advanced our understanding of human emotion experience by uncovering neural correlates of dispositional co-occurring emotions, further research is needed to elucidate the shared, common neural representations of multivariate emotion experience across individuals (Adolphs et al., 2002).

Taking the multivariate structure of human emotion into account poses a major challenge to the methodology for investigating neural representations. The representational similarity analysis (RSA) method presents a promising solution to this challenge (Kriegeskorte et al., 2008). Rather than treating each instance of multivariate emotion experience as a single value, the RSA method directly and comprehensively utilizes the multivariate data by focusing on the pairwise representational similarity distances among a group of instances and identifying the neural activity patterns that exhibit a similar distance structure with behavioral data. The RSA method has been successfully applied to explore the neural representations of various cognitive functions (Popal et al., 2019), especially for sensory processing in visual and auditory domains when the stimuli were considered to have a (hidden) multivariate property. As previously mentioned, a recent EEG study utilized an extended version of the RSA method called inter-subject RSA (Hu et al., 2022), treating each individual participant’s multivariate emotion experience across all situations as one instance. To explore the shared neural representations across individuals, a similar approach could be employed, treating the multivariate emotion experience of each situation as one instance. The pairwise similarity distances among all situations could be calculated individually and then averaged across participants. With sufficient emotional situations and participants, the participant-averaged similarity structure of situations could serve as a valid reference to identify shared neural representations.

In this study, we aimed to investigate the electrophysiological representations of multivariate human emotion experience using EEG. To elicit emotions with high ecological validity, we employed a video watching paradigm, which is popular in emotion researches. 78 participants watched 28 emotional video clips that were designed to induce emotions mainly from eight emotion categories (anger, disgust, fear, sadness, amusement, inspiration, joy, and tenderness) while their EEGs were recorded. The content of the video clips covered various emotional situations. Each video clip represented one emotional situation, and the participants rated their emotion experience on the eight emotion categories. We extracted the power spectral density (PSD) from EEG recordings as the electrophysiological representation, as suggested by previous EEG studies on emotion. As a kind of classical frequency domain analysis, the PSD represents the power distribution of EEG series in the frequency domain and has been widely utilized in EEG-based emotion recognition. An inter-situation representational similarity analysis (RSA) method was used, as described above and illustrated in Figure 1. We treated the 8-variate self-report ratings as the multivariate emotion experience and the PSD feature at each EEG recording channel in the five frequency bands (Delta, Theta, Alpha, Beta, and Gamma) as the EEG representation. The inter-situation RSA was conducted by correlating the similarity structures of situations (videos) in multivariate emotion experience and in EEG representation. Specifically, the similarity structure of situations consists of the pairwise representational dissimilarities among all situations, which were the Euclidean distances between the 8-variate ratings/PSD features (per channel, per frequency band) of the two situations. Therefore, the EEG-based electrophysiological representation of multivariate emotion experience can be identified as the PSD features with significant RSA-based correlations with the multivariate ratings. To evaluate the reliability of our findings, we conducted additional inter-situation RSA by varying the numbers of videos or participants, to test whether the identified EEG representations could be retained with a reduced number of videos/participants. Furthermore, to investigate the specificity of EEG representations for multivariate emotion experience, we conducted a series of ablation analyses by excluding each single emotion (univariate) ratings from the 8-variate ratings (leading to the corresponding 7-variate ratings), and performing the inter-situation RSA between the ablated multivariate ratings and the PSD features respectively. These analyses could explore the identified EEG representations in more detail. Additionally, the EEG representations of univariate emotion experience were also explored by applying the inter-situation RSA. As probably the first study in this direction, the present study is exploratory in nature without prior hypotheses. The findings are expected to extend our understanding of the neural mechanisms of human emotion in the multivariate context.

**Figure 1.**
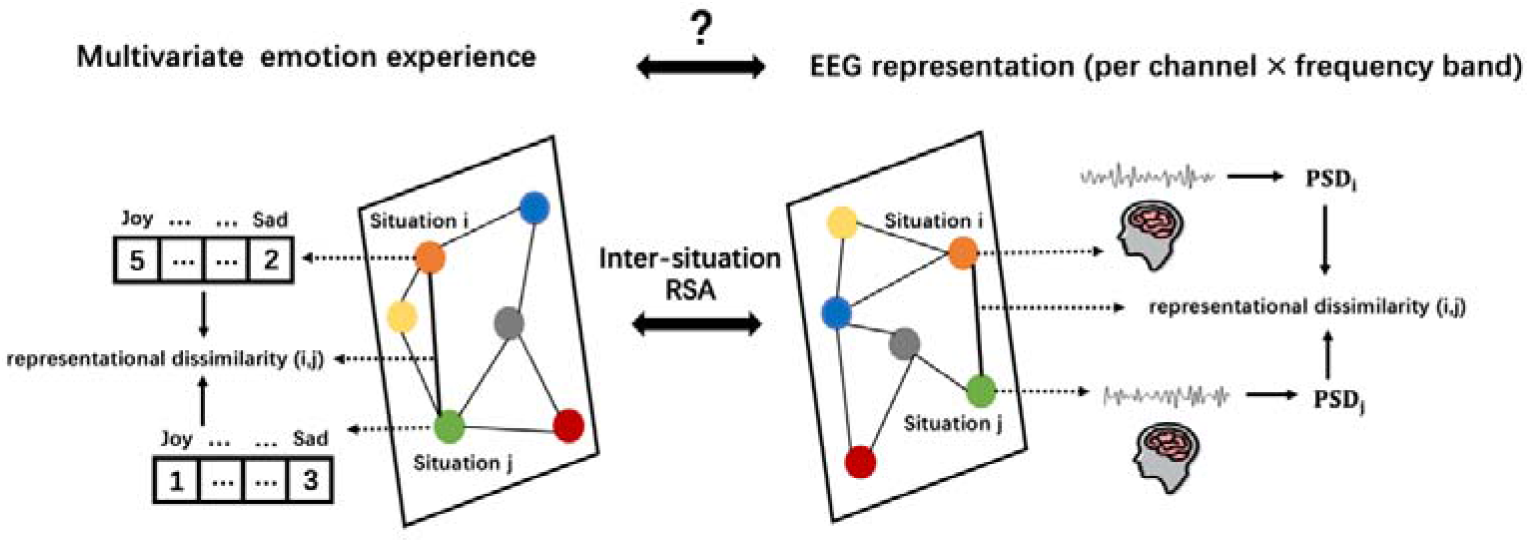
Schematic diagram of inter-situation representational similarity analysis (RSA). The inter-situation RSA is conducted based on the similarity structures of situations in multivariate emotion experience and in EEG representation (per channel, per frequency band). The similarity structure of situations consists of the pairwise representational dissimilarities among all situations. As a simplified example, the representational dissimilarities between Situation i and Situation j in multivariate emotion experience and in EEG representation, can be calculated by the Euclidean distances between their 8-variate emotion experience ratings and between their power spectral density (PSD) features at one of the 30 EEG recording channels, in one of the five frequency bands (Delta, Theta, Alpha, Beta, and Gamma), respectively.

## Method

### The dataset

As part of a larger study on emotion, the present study used a dataset with 78 participants watching 28 emotional video clips. The sample size was determined empirically with reference to recent EEG studies on emotion. The details of the dataset^2^ are introduced as below.

78 Chinese college students in Beijing were recruited (30 males, 48 females, aged 17–24, mean age 20.2 years). All participants were right-handed, with normal or corrected-to-normal vision and audition. The study was approved by the local Ethics Committee of Tsinghua University. Written informed consent was obtained from all participants.

Twenty-eight video clips selected from previous published emotional video databases were used (Hu et al., 2022). Among these videos, 12 videos were used primarily to elicit four negative emotions (anger, disgust, fear, and sadness), with three videos targeting at one of the emotion categories; another 12 videos were used primarily to elicit four positive emotions (amusement, inspiration, joy, and tenderness), again with three videos targeting at one of the emotion categories; there were additionally four videos for eliciting neutral emotion experience. The duration of these videos varied between 34 s to 129 s, with a mean value of 63 s. Chinese subtitles were provided for all videos to ensure the participants’ understanding of the video content. More detailed information on the video clips could be found in the Supplementary Materials, Table S1.

The experiment was conducted in a laboratory environment with normal ambient light. Participants were seated about 60 cm away from a 22-inch LCD monitor (Dell, USA), and the sound was played by stereo speakers (Dell, USA). These videos were presented in seven blocks. To reduce the possible influence of alternating valence, each block contained four trials with the same valence videos, resulting in three negative blocks, three positive blocks, and one neutral block. In each trial, the participants were first presented with a 5-second fixation cross in the center of the screen, followed by the presentation of a video. After the completion of each video presentation, the participants rated their emotion experience on 7-point Likert scales on the eight emotion categories (anger, disgust, fear, sadness, amusement, inspiration, joy, and tenderness). The participants then took a rest for at least 30 s, and decided when start to watch the next video clip by keypress. Between two successive blocks, the participants were asked to complete 20 arithmetic questions (the distraction task), in order to minimize the influence of previously elicited emotion experience on the next block.

A portable wireless EEG system (NeuSen.W32, Neuracle, China) was used to record EEG signals from 32 channels placed according to the international 10-20 system: Fp1/2, Fz, F3/4, F7/8, FC1/2, FC5/6, Cz, C3/4, T3/4, CP1/2, CP5/6, T5/6, Pz, P3/4, PO3/4, Oz, O1/2, A1/2 (left and right mastoids). Channels were referenced to CPz with a forehead ground at AFz. The sampling rate was 250 Hz. Channel impedances were kept below 10 kOhm for all channels.

### Behavioral data analysis

As previously mentioned, each video in the present study was designed to represent an emotional situation, and the ratings on all 8 emotion categories for each video were used as the multivariate (8-variate) emotion experience of that situation. These 8 emotion categories were chosen to reflect the relatively rich and complex nature of human emotion.

### EEG data analysis

The raw EEG signals were first re-referenced to the average of the mastoids (A1/2). Then, a two-pass fourth-order Butterworth filter was applied to bandpass filter the signals to 0.05-47 Hz. To remove artifacts related to eye and muscle movements, independent component analysis (ICA) was used, with approximately 1-2 artifact-related independent components (ICs) removed per participant. To capture the maximal emotional responses and match with the final emotional states (that is the multivariate emotion experience ratings), the EEG data corresponding to the last 30 seconds of each video were extracted, following previous studies (Koelstra et al., 2011). All EEG data pre-processing was conducted in MATLAB, using the FieldTrip Toolbox (Oostenveld et al., 2011).

The EEG features in this study were defined as the power spectral density (PSD) per channel (30 EEG channels) per frequency band (Delta: 1-3 Hz, Theta: 3-8 Hz, Alpha: 8-14 Hz, Beta: 14-30 Hz, and Gamma: 30-47 Hz), resulting in 150 PSD features (30 channels × 5 frequency bands). These features of each participant were calculated from the 30-second cleaned EEG recordings of each video, by computing the Fourier Transform-based PSD (NFFT = 256, the length of Hamming window = 125, and overlap = 50%) over the five frequency bands for each EEG channel. Finally, the resulting EEG data were 78 (participants) × 28 (videos) × 150 (PSD features).

### Inter-situation representational similarity analysis

In order to investigate how the multivariate (co-occurring) emotions are represented in EEG activity during video watching, we conducted an inter-situation representational similarity analysis (RSA). This involved measuring the correlation between the similarity structure of the multivariate emotion experience ratings and that of the PSD feature per channel per frequency band. The similarity structure was represented by a dissimilarity matrix, which was calculated as the Euclidean distance between the representational vectors of each video (situation) pairs (the vector dimensions for ratings and PSD feature are 8 and 1, respectively). We aimed to reduce individual differences and capture shared representations across individuals by calculating dissimilarity matrices for each participant and then averaging them across all participants. This yielded a series of participant-averaged 28 × 28 dissimilarity matrices. We then performed Spearman correlations between the upper triangles of the dissimilarity matrix of multivariate emotion experience ratings and each of the 150 dissimilarity matrices of PSD features, respectively. To assess the statistical significance of each correlation value, we conducted a permutation test by shuffling the video order and recalculating the correlations 1000 times to form a null distribution. The true correlation value was compared with the null distribution to obtain the p value. All the obtained p values were corrected for multiple comparisons using the false discovery rate (FDR) (Storey et al., 2002), with a significant level as 0.05. The EEG representations for multivariate emotion experience are reflected as the PSD features with significant correlations. The inter-situation RSA calculation process is illustrated in the Supplementary Materials, Figure S1.

To assess the reliability of the EEG representations, we conducted additional inter-situation RSA by varying the numbers of videos or participants. Firstly, we performed inter-situation RSA with 9 or 18 videos, where one (or two) video(s) targeting at each of the nine emotion categories (four negative, four positive and neutral) were randomly sampled to create 9-video (or 18-video) conditions. We generated 1000 different random sampling cases for each condition and performed inter-situation RSA in each condition, to examine whether or how the EEG representation could be retained when the numbers of videos (situations) were reduced from 28 to 18 and 9. Secondly, we conducted inter-situation RSA with another four conditions with the numbers of participants as 10, 20, 40, and 60, respectively. In each condition, we randomly shuffled the participant order 1000 times and selected the top N participants (N = 10, 20, 40, or 60) in each random shuffling, creating 1000 different sampling cases. Similarly, we performed inter-situation RSA in each condition, to examine the reliability of EEG representation with a reduced number of participants.

Furthermore, to investigate the specificity of EEG representation for multivariate emotion experience, we conducted a series of ablation analyses: we excluded each univariate ratings from the 8-variate ratings respectively, leading to the corresponding 7-variate ratings, and then conducted the inter-situation RSA between these ablated multivariate ratings and the 150 PSD features, respectively. These analyses could demonstrate the impact of each emotion experience variate (emotion category) on the identified EEG representation. In addition, we also investigated the EEG representation of univariate emotion experience, by conducting the inter-situation RSA between each univariate ratings and the 150 PSD features respectively. The above analyses could indicate whether the multivariate emotion experience has the specific EEG representation which is different with that of the univariate emotion experience.

## Result

In the behavioral data analysis, all videos have successfully elicited the intended emotion experience: all the negative and positive videos rated highest on their target emotion categories (mean ratings per target emotion category: anger 5.03 ± 1.84, disgust 5.13 ± 1.97, fear 4.60 ± 1.99, sadness 4.71 ± 1.61, amusement 5.03 ± 1.61, inspiration 5.21 ± 1.67, joy 4.41 ± 1.51, tenderness 5.40 ± 1.24), and the neutral videos evoked emotion experience with the lowest intensity. The cross-participants averaged ratings of the eight emotion categories for all 28 videos are presented in the Supplementary Materials, Figure S2. The effectiveness of emotion elicitation is further suggested by the overall significant differences from a series of paired t-tests, showing that the ratings of one specific emotion category in its targeting videos (the videos with this emotion category as the intended elicitation target) were significantly higher than that in any other videos. More details are revealed in the Supplementary Materials, Table S2. Notably, the 28 videos served as relatively rich emotional situations and the induced emotion experience was not a single emotion but rather a co-occurrence of multiple emotions (e.g., participants experienced anger as well as disgust and sadness while watching anger videos), supporting our proposed view of a multivariate emotion experience.

The EEG-based electrophysiological representations of the multivariate emotion experience using inter-situation representational similarity analysis (RSA) are illustrated in Figure 2a. Out of the 150 PSD features, 70 showed significant correlations with the multivariate emotion experience ratings, distributed over 1 channel in the Delta band, 6 channels in the Theta band, 29 channels in the Alpha band, 21 channels in the Beta band, and 13 channels in the Gamma band. The channels with the highest RSA correlation value (r value) in each frequency band were the Fp2 in the Delta band (r = 0.2434, p < 0.001), the Fp2 in the Theta band (r = 0.2067, p < 0.001), the O2 in the Alpha band (r = 0.3286, p < 0.001), the PO4 in the Beta band (r = 0.3386, p < 0.001), and the T5 in the Gamma band (r = 0.2301, p < 0.001). As the significant correlations were mainly observed in the Alpha and Beta bands, the PSD feature at O2 in the Alpha band, and PSD feature at PO4 in the Beta band were chosen as representative examples to show how the EEG representations were associated with the multivariate emotion experience ratings, as shown in Figure 2b.

**Figure 2.**
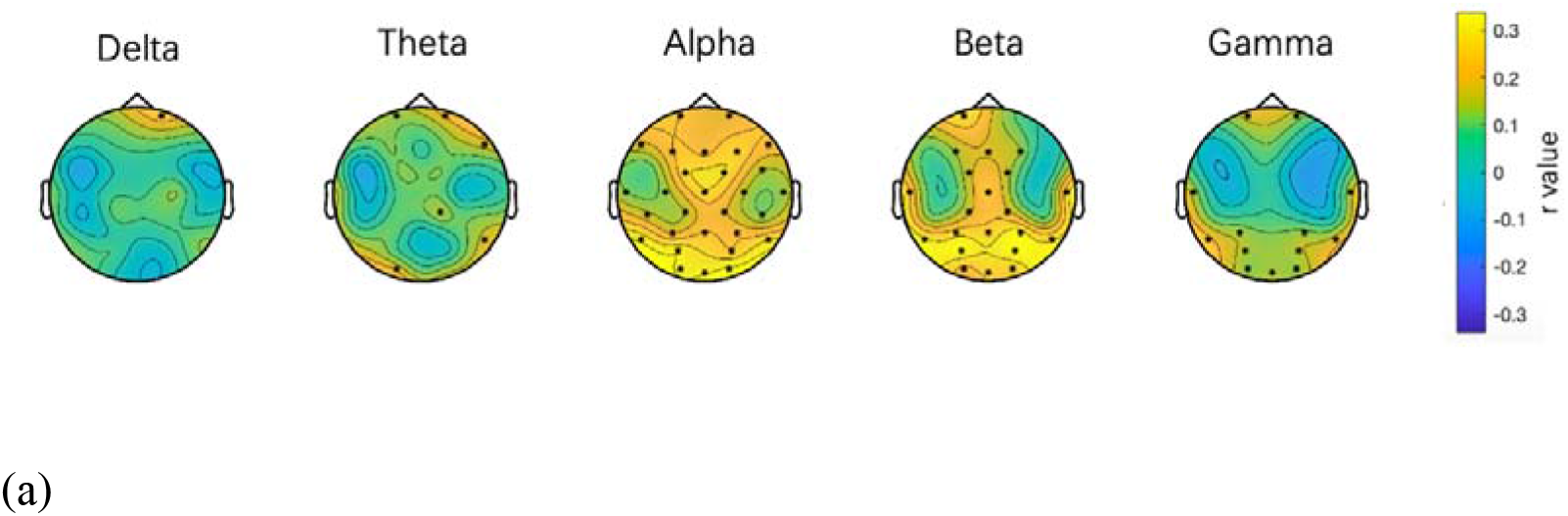
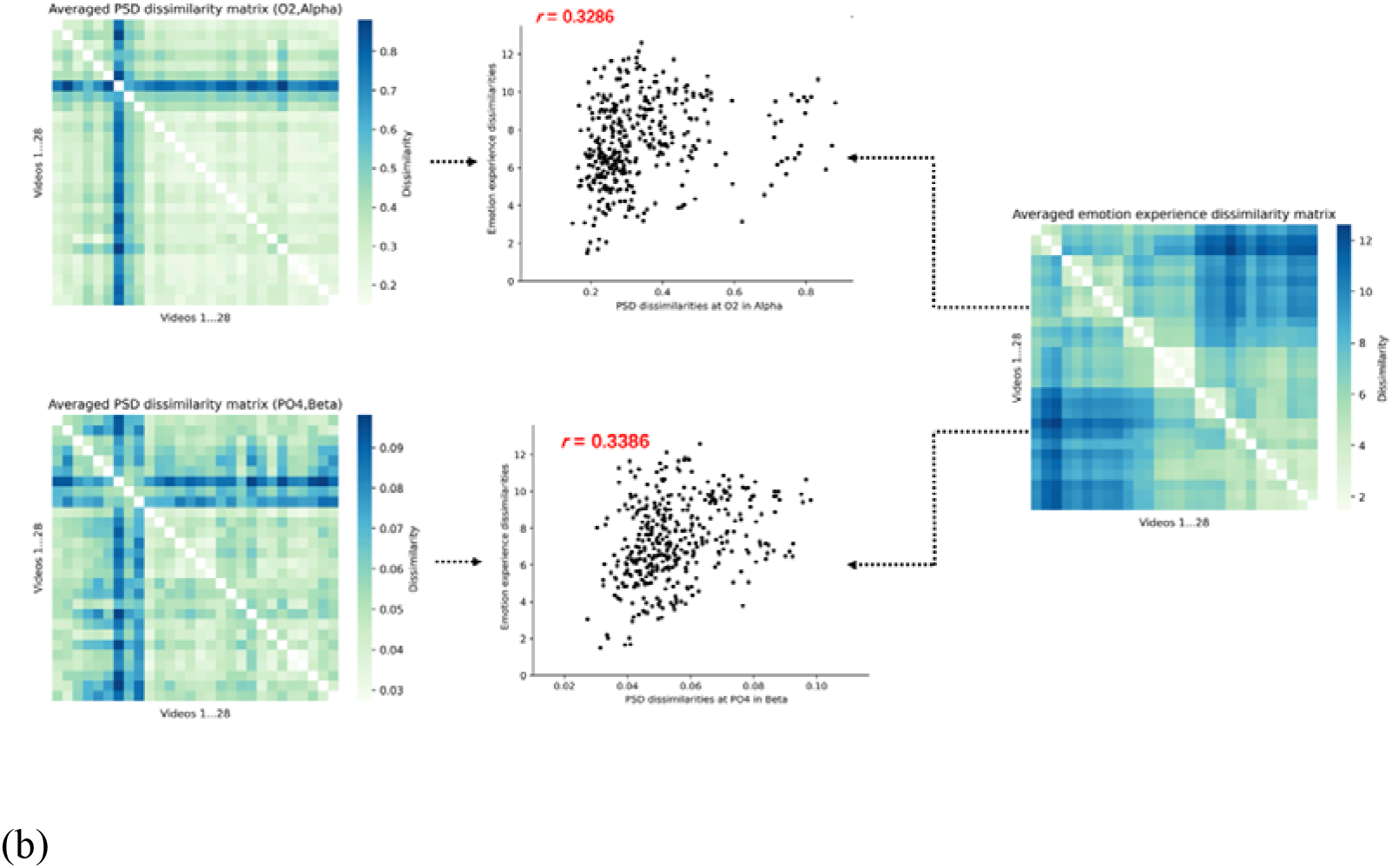
(a) The inter-situation RSA correlation values between multivariate emotion experience ratings and the PSD feature at each channel in the five frequency bands. The channels with significant correlation values (ps < 0.05, FDR corrected) are marked in the topographies. (b) The participant-averaged dissimilarity matrices of two representative PSD features (O2 in the Alpha band, and PO4 in the Beta band) are shown in left, and the participant-averaged dissimilarity matrix of multivariate emotion experience ratings is shown in right. The scatter plots present the pairwise dissimilarities among all videos.

The investigations on the reliability of the EEG-based electrophysiological representations of the multivariate emotion experience are summarized in the Supplementary Materials, Figure S3. We took the PSD feature at PO4 in the Beta band as one example (showing the highest inter-situation RSA correlation value, r = 0.3386), to show the RSA-based correlation values (r values) in all the 1000 random sampling cases of each video/participant condition. It can be seen in Figure S3a that the r values did not decrease much when the numbers of videos (situations) were reduced to 18 and 9 or when there were more than 40 participants: the median r values were 0.3144 (p < 0.001) and 0.2946 (p = 0.071) for 18-video and 9-video conditions, respectively; the median r values were 0.3003 (p < 0.001), 0.2641 (p < 0.001), 0.2109 (p < 0.001), and 0.1601 (p < 0.05) for the 60-participant, 40-participant, 20-participant and 10-participant conditions, respectively. Furthermore, we selected the cases corresponding to these median r values as the representative cases, to show the inter-situation RSA correlation values of all the 150 PSD features in the representative case of each video/participant condition. In general, the EEG representations of multivariate emotion experience were largely preserved with more than 18 videos (see Figure S3b) or 40 participants (see Figure S3c), although the number of significant correlations decreased.

The specificity of EEG-based electrophysiological representations for the multivariate emotion experience is illustrated in the Supplementary Materials, Figure S4. By excluding each univariate ratings from the 8-variate ratings respectively, the RSA-based ablation analyses revealed that the EEG representations of multivariate emotion experience were relatively stable and consistent, suggesting a limited impact of single emotion experience variate (see Figure S4a). The EEG representations of univariate emotion experience are shown in Figure S4b, which were different with those of multivariate emotion experience and were varied among emotion categories: negative univariate emotion experience (especially disgust and fear) generally had more significant EEG representations than positive univariate emotion experience.

## Discussion

This study aimed to investigate the EEG-based electrophysiological representation of human emotions by addressing the complex and co-occurring nature with multivariate emotion construct. To achieve this, the study explored the association between the multivariate emotion experience during video watching (self-report ratings on 8 emotion categories), and the power spectral density (PSD) features at each EEG recording channel in the five frequency bands (Delta, Theta, Alpha, Beta, and Gamma), by using the inter-situation representational similarity analysis (RSA). Results revealed that the brain activities in Alpha and Beta bands, mainly over the frontal and parietal-occipital regions were closely associated with the multivariate emotion experience, which was found to be reliable with sufficient situations and participants. Moreover, the multivariate emotion experience was found to have stable and specific EEG representations, which were different from those univariate-based findings. These findings demonstrate that taking a multivariate perspective could facilitate our understanding of neural mechanisms of human emotions.

In the behavioral data analysis, it is interesting to find that the emotional situations targeting one specific emotion category could evoke same-valence emotions as well as some opposite-valence emotions (e.g., participants reported that they felt some tenderness when watching sadness videos). In view of this, the traditional perspective of univariate emotion experience may be insufficient to describe the holistic picture of real emotion experience (in other words, it is incomplete), and the multivariate structure of human emotion could be alternative and its validity has been proved in both laboratory setting (Larsen et al., 2014) and daily life.

The findings of a widely-distributed, Alpha-and-Beta-dominated EEG-based electrophysiological representations of the multivariate emotion experience are consistent and extend the existing literatures mainly with single emotions. Firstly, the association between these brain oscillations and emotion processing has been consistently reported in previous EEG studies. Specifically, the functional roles of Alpha and Beta activities have been widely acknowledged, for emotion regulation, emotional memory processing, etc. Accordingly, the EEG features extracted from the Alpha and Beta bands also achieved competent performance on emotion recognition tasks (Wang et al., 2011) in the affective computing field. Secondly, the identified widely-distributed brain regions including frontal, central, and parietal-occipital areas in this study are in accordance with the emerging network view on emotion processing. While different brain regions have been attributed to different emotion categories (e.g., disgust, fear, sadness, happiness) (Hamann et al., 2012), the present multivariate results showed a comprehensive coverage of all these brain regions. Moreover, given the overall inclusion of multiple emotion categories (emotion experience variates) in the present study, the observed neural activities over the prefrontal and midline regions could be considered as important further evidence for their functional roles for general-purpose emotion processing, such as emotional salience processing, self-relevant and introspective processing, as well as integration of internal, mental, and bodily states. The occipital representations, however, could be related to visual cortex activation by video presentation, or alternatively the emotion perception of visual stimuli (Junghöfer et al., 2001). Taken together, whereas the electrophysiological representations of multivariate emotion experience are compatible with the univariate-based findings, the present findings provide an overall description of the neural activities related to emotion processing in general from a multivariate perspective.

The reliability analyses provided an important support for the stability of the revealed electrophysiological representations. It was found that the identified EEG representations based on 28 videos and 78 participants were largely remained, when the numbers of videos or participants were reduced to 18 or 40, respectively. Hereby, the representations could be considered as a robust finding given a sufficient number of emotional situations and participants. Furthermore, the ablation analyses illustrated the stability of the representations when excluding each of the single emotion category, even if some of these emotion categories were associated with significant neural responses in univariate analysis (see Figure S4b, such as disgust and fear). Taken together, we demonstrated that the present findings were not dependent on specific situations, participants, or emotion categories. Therefore, it is plausible to consider the observed representations for characterizing a general multivariate emotion processing shared across participants. Considering the previous study has observed the distinct neural representation when the emotion blended or not (Zinner et al., 2008), it is interesting to conduct a deeper investigation to explore the general and stable neural representation reflecting the co-occurring nature of human emotions.

It is worth noting that the present findings together with a recent individual difference study provide complementary information regarding the neural representation of multivariate emotion experience. In that study, researchers employed inter-subject RSA method and reported significant correlations between inter-participant similarity of multivariate emotion experience and Delta-band EEG activities over prefrontal and temporo-parietal regions, and Theta-band activities over frontal regions (Hu et al., 2022). The shared neural representations across participants reported in the present study seemed to involve overlapped (prefrontal and frontal) but distinct (no temporal but additionally occipital) brain regions, suggesting a possible differentiation of brain regions for representing individualized and shared emotion processing. The possible distinct representations for these two types of emotion processing were further supported by the different frequency bands: whereas the Delta and Theta bands were shown to be involved for individual difference, the Alpha and Beta bands were mainly responsible for shared processing. The distinction echoed studies in other fields, for example, the previous study (Seghier et al., 2016) showed the whole-brain consistency and variability in semantic-related brain responses across participants, indicating that the inferences at the group level were not always relevant (or valid) at the individual level. This could be interpreted by the hypothesis that each individual brain-activation map was a sum of two quantities: the mean group effect plus an individual-specific effect (Seghier et al., 2018). The mean group effect (the central tendency) could illustrate the similar activation pattern of core functional networks across participants, while the inter-subject variability (the deviation from the group) may reflect the individual differences in cognitive style and preferred cognitive strategy, as well as the alternative neural systems for a given task, including motor, visual, semantic, default mode and oculomotor networks, etc. Hence, the identified general representation of multivariate emotion processing in the current study, could be an important complement and enrich the conclusions that have been drawn from the individual-specific perspective.

There are several limitations that should be noted. Firstly, although EEG is advantageous for its high temporal resolution and rich spectral information, high ecological validity, its relatively low spatial resolution has limited our exploration about the brain regions involved for multivariate emotion experience. Neuroimaging techniques with a high spatial resolution such as fMRI are necessary to provide further information in this regard, and the inter-situation RSA method could be readily applicable for such analyses. Secondly, while the present study followed the majority of recent emotion studies to have videos as the emotion elicitation materials, the generalizability of the present findings needs to be further verified with more types of emotion tasks, such as music listening, emotion experience recall, etc. Moreover, the complexity of the emotion elicitation materials could be further improved: while the present materials showed sufficient richness of emotion experience within the basic emotions, further studies could include materials with simultaneous elicitation of the high-level emotions just like aesthetic emotions, in order to have a more comprehensive overview of the neural representations of the multivariate emotion experience.

## Disclosure of interest

The authors report no conflict of interest.

1. In the present study, we use the ‘multivariate emotion experience’ to represent the co-occurring pattern across multiple emotion categories, and the ‘univariate emotion experience’ corresponds to the emotion experience within a single emotion category. Specifically, each emotion category is considered as one variate.
2. The data are available via https://osf.io/cfygx/. This study’s design and its analysis were not preregistered.

