## supplemental file for "Electrophysiological Representations of Multivariate Human Emotion Experience"

**Supplementary Materials**

**Table S1.** *Information about the video clips*

| **Clip Number** | **Source Film** | **Targeted Emotion** | **Source Database** | **Duration（s）** | **Language** |
| --- | --- | --- | --- | --- | --- |
| 1 | The Tokyo Trial | Anger | DCEF | 81 | Japanese\Chinese |
| 2 | Documentary about the Nanjing Massacre | Anger | \ | 63 | English |
| 3 | City of Life and Death | Anger | DCEF | 73 | \ |
| 4 | Trainspotting | Disgust | FilmStim | 78 | English |
| 5 | Indiana Jones and the Last Crusade | Disgust | FilmStim | 69 | English |
| 6 | Hellraiser | Disgust | FilmStim | 90 | English |
| 7 | The Shining | Fear | FilmStim | 56 | English |
| 8 | The Shining | Fear | \ | 60 | English |
| 9 | The Exorcist | Fear | FilmStim | 105 | English |
| 10 | In Bruges | Sadness | PED | 45 | English |
| 11 | Departures | Sadness | PED | 60 | Japanese |
| 12 | Gangs of New York | Sadness | PED | 81 | English |
| 13 | Blue | Neutral | FilmStim | 35 | English |
| 14 | Blue | Neutral | FilmStim | 44 | English |
| 15 | Blue | Neutral | FilmStim | 38 | English |
| 16 | Blue | Neutral | FilmStim | 43 | English |
| 17 | Modern Times | Amusement | PED | 55 | English |
| 18 | Minions | Amusement | PED | 69 | English |
| 19 | Mr. Bean | Amusement | PED | 73 | English |
| 20 | Forrest Gump | Inspiration | PED | 129 | English |
| 21 | The Theory of Everything | Inspiration | PED | 77 | English |
| 22 | The Shawshank Redemption | Inspiration | PED | 83 | English |
| 23 | My Neighbor Totoro | Joy | PED | 34 | Japanese |
| 24 | Night at the Museum Ⅲ | Joy | PED | 37 | English |
| 25 | Harry Potter Ⅰ | Joy | PED | 67 | English |
| 26 | The Pursuit of Happiness | Tenderness | PED | 63 | English |
| 27 | Juno | Tenderness | PED | 83 | English |
| 28 | Sex and the City Ⅱ | Tenderness | PED | 77 | English |

*Note*: DCEF refers to the standardised database of Chinese emotional film clips (Ge et al., 2019), PED refers to the positive emotion database (Hu et al., 2019), and FlimStim refers to the database created by Schaefer (2010).

**Table S2.** *Self-Reported Emotion Experience Ratings, Means and Standard Deviations by Video Targeting Emotion Category*

| Self-Reported Emotion Category | Video Targeting Emotion Category | | | | | | | | |
| --- | --- | --- | --- | --- | --- | --- | --- | --- | --- |
|  | Anger | Disgust | Fear | Sadness | Neutral | Amusement | Inspiration | Joy | Tenderness |
| Anger | 5.03 (1.84) | 0.61 (1.31)  *t* = -30.27* | 1.09 (1.71)  *t* = -24.23* | 0.86 (1.35)  *t* = -28.22* | 0.22 (0.61)  *t* = -43.59* | 0.41 (0.83)  *t* = -35.36* | 0.26 (0.62)  *t* = -38.01* | 0.23 (0.64)  *t* = -38.05* | 0.21 (0.58)  *t* = -38.65* |
| Disgust | 2.74 (2.42)  *t* = -11.81* | 5.13 (1.97) | 2.47(2.29) *t* = -13.59* | 0.69 (1.33)  *t* = -28.82* | 0.19 (0.56)  *t* = -42.56* | 0.26 (0.63)  *t* = -36.40* | 0.66 (1.35)  *t* = -28.88* | 0.18 (0.55)  *t* = -37.45* | 0.18 (0.52)  *t* = -37.53* |
| Fear | 1.79 (2.06)  *t* = -15.16* | 2.92 (2.33)  *t* = -8.47* | 4.60 (1.99) | 1.01 (1.54)  *t* = -22.08* | 0.26 (0.69)  *t* = -36.20* | 0.21 (0.57)  *t* = -32.78* | 0.34 (0.83)  *t* = -30.57* | 0.24 (0.67)  *t* = -32.16* | 0.20 (0.57)  *t* = -32.88* |
| Sadness | 4.27 (2.26)  *t* = -2.39* | 0.99 (1.56)  *t* = -25.48* | 1.75 (2.03)  *t* = -17.59* | 4.71 (1.61) | 0.74 (1.36)  *t* = -31.41* | 0.37 (0.82)  *t* = -36.93* | 0.70 (1.26)  *t* = -30.23* | 0.18 (0.57)  *t* = -40.92* | 0.75 (1.28)  *t* = -29.60* |
| Amusement | 0.10 (0.22)  *t* = -46.90* | 0.67 (1.23)  *t* = -33.26* | 0.28 (0.76)  *t* = -41.36* | 0.12 (0.23)  *t* = -46.71* | 0.36 (0.82)  *t* = -44.73* | 5.03 (1.61) | 1.30 (1.82)  *t* = -23.79* | 1.74 (1.77)  *t* = -21.26* | 0.96 (1.41)  *t* = -29.51* |
| Inspiration | 0.53 (1.26)  *t* = -34.53* | 0.28 (0.63)  *t* = -42.57* | 0.25 (0.61)  *t* = -42.90* | 0.60 (1.08)  *t* = -35.71* | 0.76 (1.18)  *t* = -36.80* | 1.44 (1.64)  *t* = -24.80* | 5.21 (1.67) | 2.47 (1.98)  *t* = -16.30* | 3.12 (2.01)  *t* = -12.30* |
| Joy | 0.17 (0.33)  *t* = -42.30* | 0.46 (0.86)  *t* = -35.06* | 0.29 (0.69)  *t* = -38.23* | 0.27 (0.49)  *t* = -40.10* | 1.43 (1.54)  *t* = -22.76* | 4.33 (1.57)  *t* = -0.48 | 3.98 (1.70)  *t* = -2.88* | 4.41 (1.51) | 4.17 (1.47)  *t* = -1.68 |
| Tenderness | 0.18 (0.41)  *t* = -61.67* | 0.33 (0.67)  *t* = -55.61* | 0.33 (0.70)  *t* = -55.06* | 2.02 (2.07)  *t* = -21.70* | 1.18 (1.51)  *t* = -35.24* | 1.92 (1.79)  *t* = -24.70* | 3.15 (2.11)  *t* = -14.23* | 3.49 (2.03)  *t* = -12.40* | 5.40 (1.24) |

*Note.* The means of self-reported ratings on each emotion category by video targeting emotion category are shown with the corresponding standard deviations in brackets. The *t*-values are from paired *t*-tests where the ratings of each participant were averaged across the videos targeting the same emotion category. The paired *t*-tests contrasted the averaged ratings of one specific emotion category across its targeting videos (the videos with this emotion category as the intended elicitation target) with that across other videos (e.g., averaged ratings of Anger across the videos targeting Anger versus the videos targeting Disgust revealed a significant difference with *t* = -30.27). Significant results (*p*s<0.05, FDR corrected) are marked with asterisk.

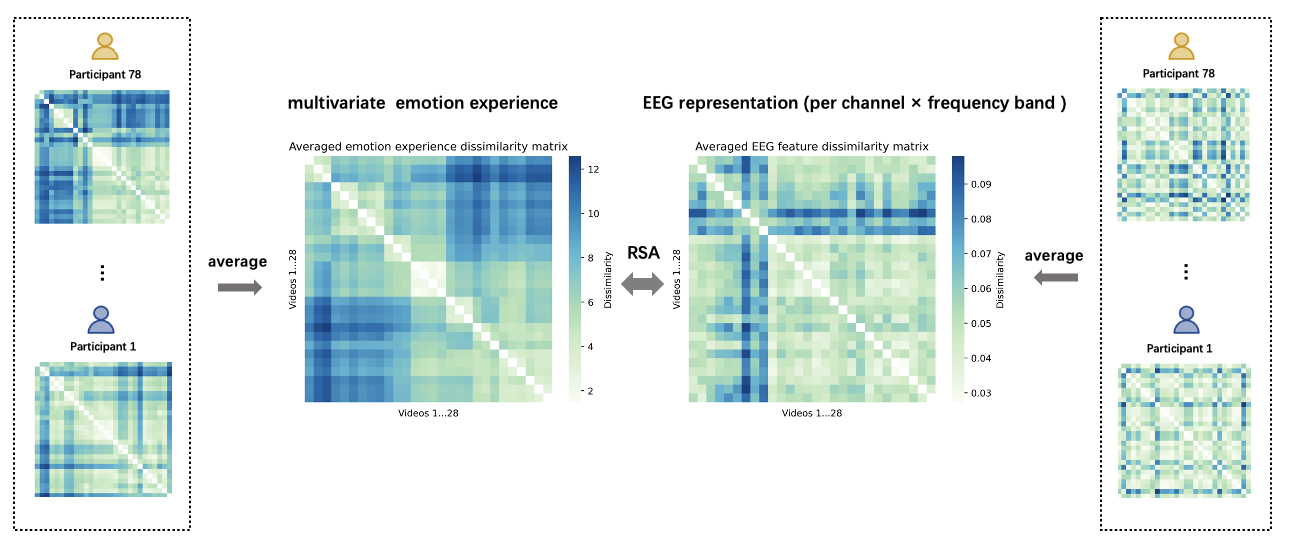

**Figure S1.** The calculation process of inter-situation representational similarity analysis (RSA). The similarity structures of videos in multivariate emotion experience and in EEG representation were presented by their dissimilarity matrices respectively.

Note: EEG representation is the power spectral density (PSD) feature at one of the 30 EEG recording channels, in one of the five frequency bands (Delta, Theta, Alpha, Beta, and Gamma).

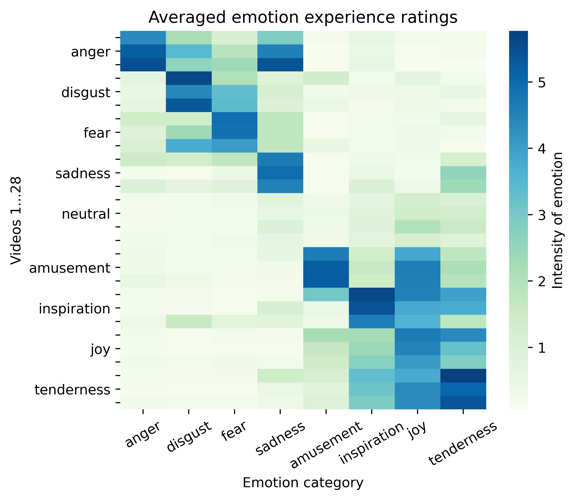

**Figure S2.** The participant-averaged ratings of each video on eight emotion categories. The videos were arranged based on their target emotion categories: videos 1-3, 4-6, 7-9, 10-12, 13-16, 17-19, 20-22, 23-25, 26-28 targeted the emotion categories of anger, disgust, fear, sadness, neutral, amusement, inspiration, joy and tenderness, respectively.

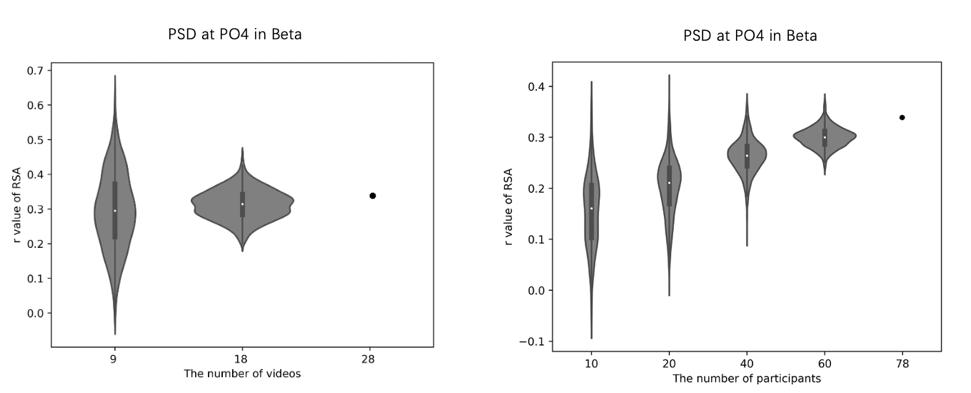

(a)

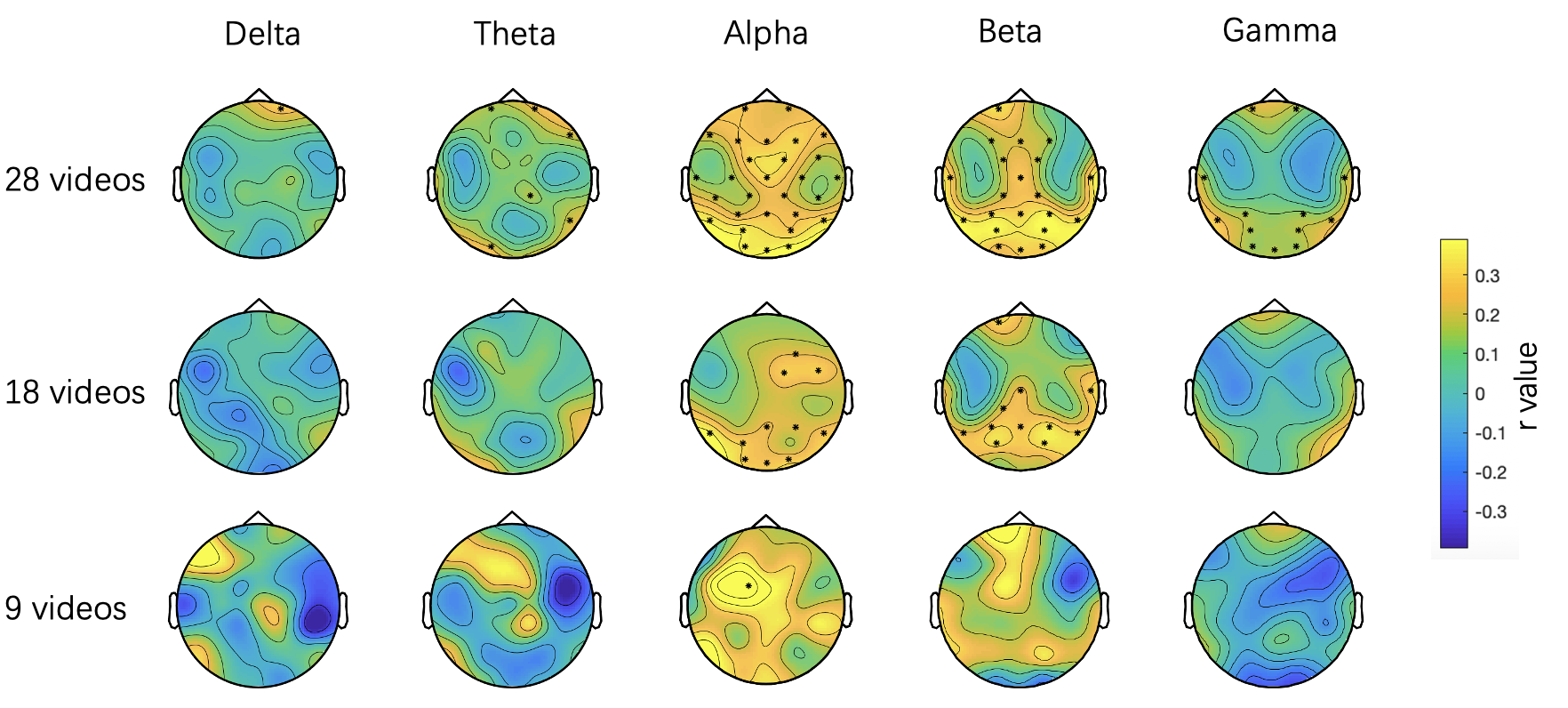

(b)

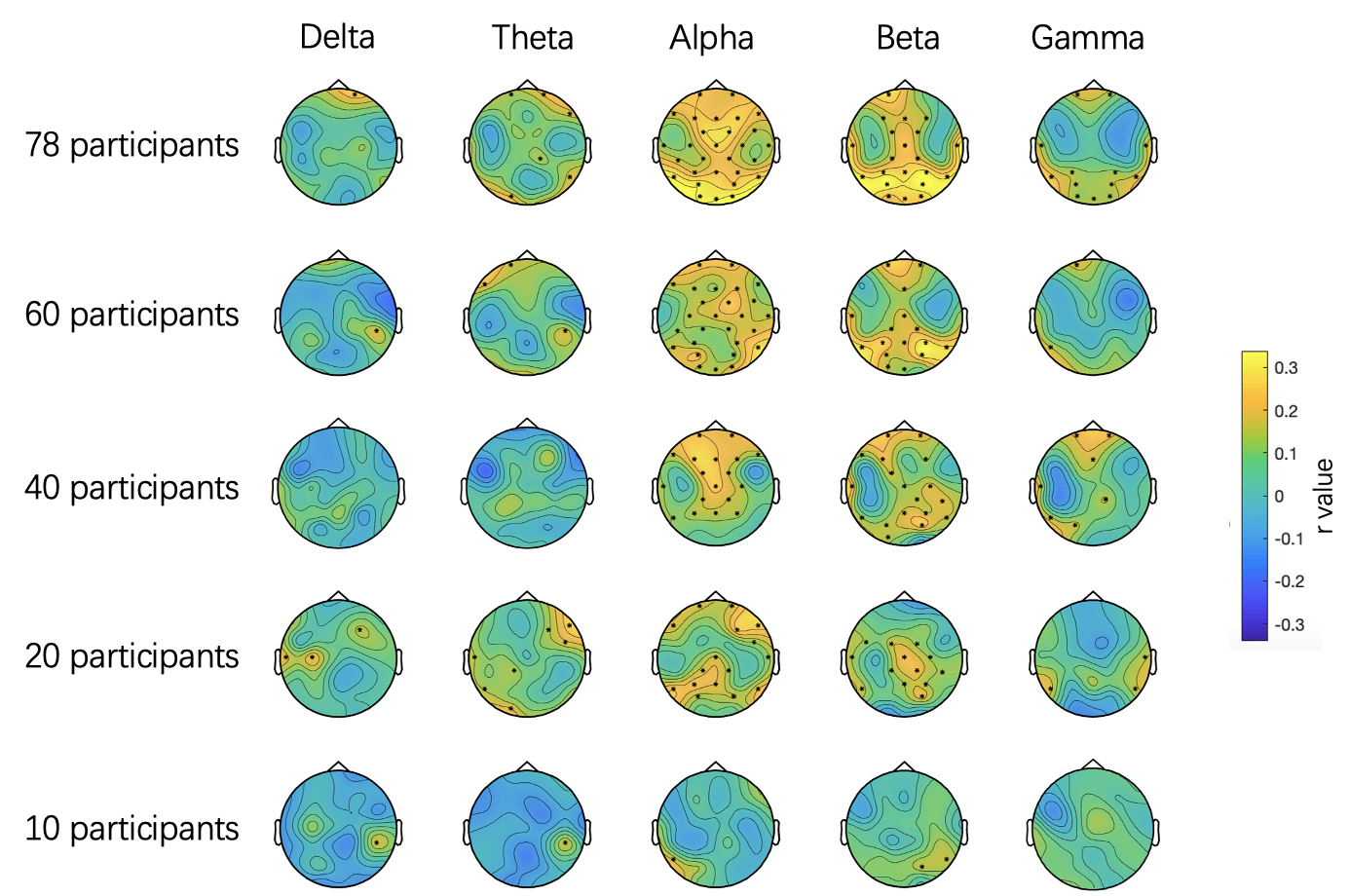

(c)

**Figure S3.** The inter-situation RSA correlation values (r values) between multivariate emotion experience ratings and the PSD features, in three video conditions (9-video, 18-video, and 28-video) and five participant conditions (10-participant, 20-participant, 40-participant, 60-participant, and 78-participant). (a) Taking the PSD feature (PO4, Beta) as one example, its r values in all the 1000 random sampling cases of each video condition are shown in the left, and of each participant condition are shown in the right. The r values of the whole PSD features (at each channel in the five frequency bands) in the representative case of each video/participant condition, are shown in (b) and (c) respectively. The channels with significant correlation values (ps < 0.05, FDR corrected) are marked in the topographies.

Note: Among all the 1000 cases of each condition, the one with median r value of the PSD feature (PO4, Beta) was selected as a representative case for the condition. The topographies corresponding to 28-video condition in (b) and the topographies corresponding to 78-participant condition in (c) are the same as Figure 2a.

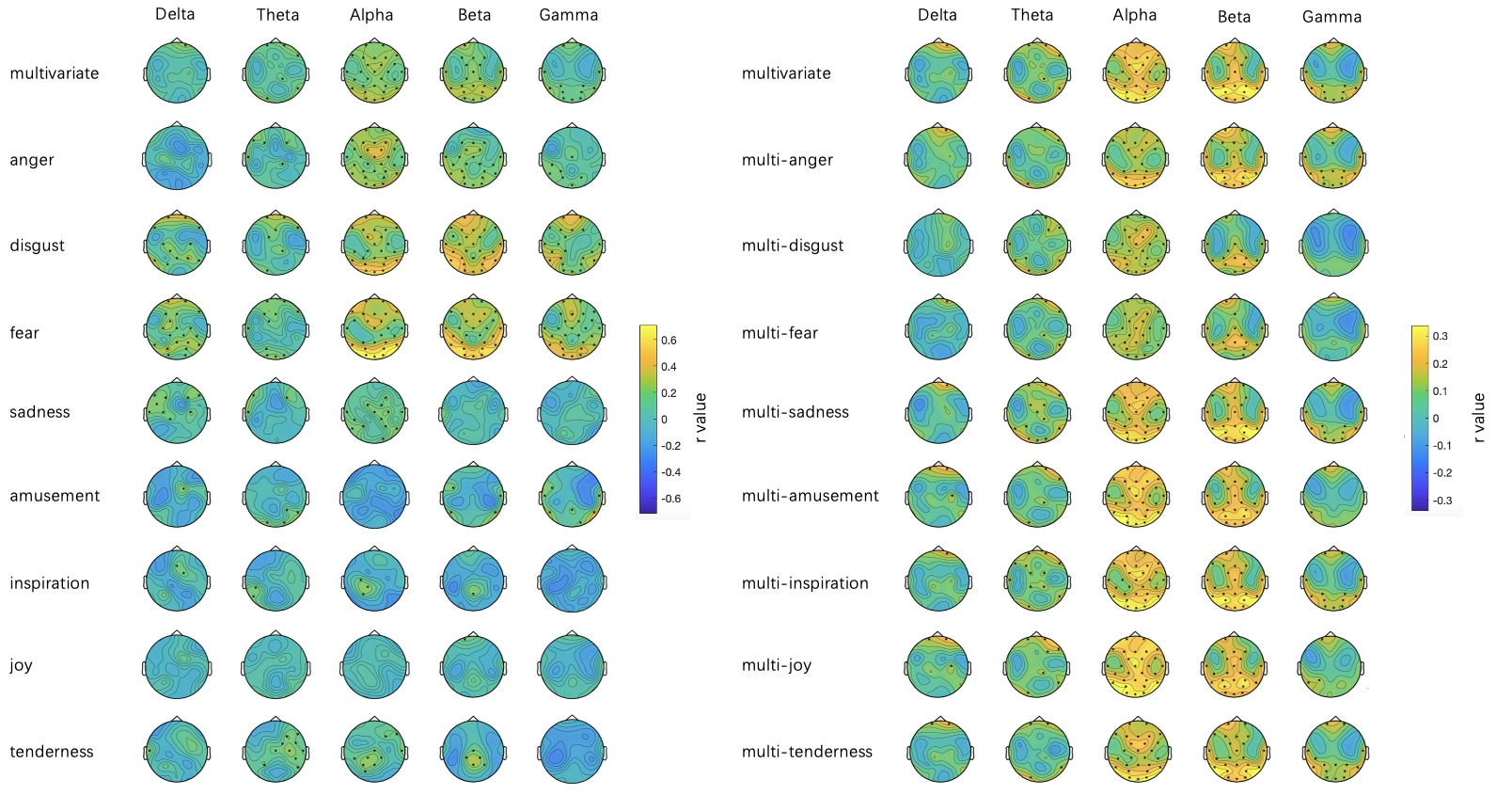

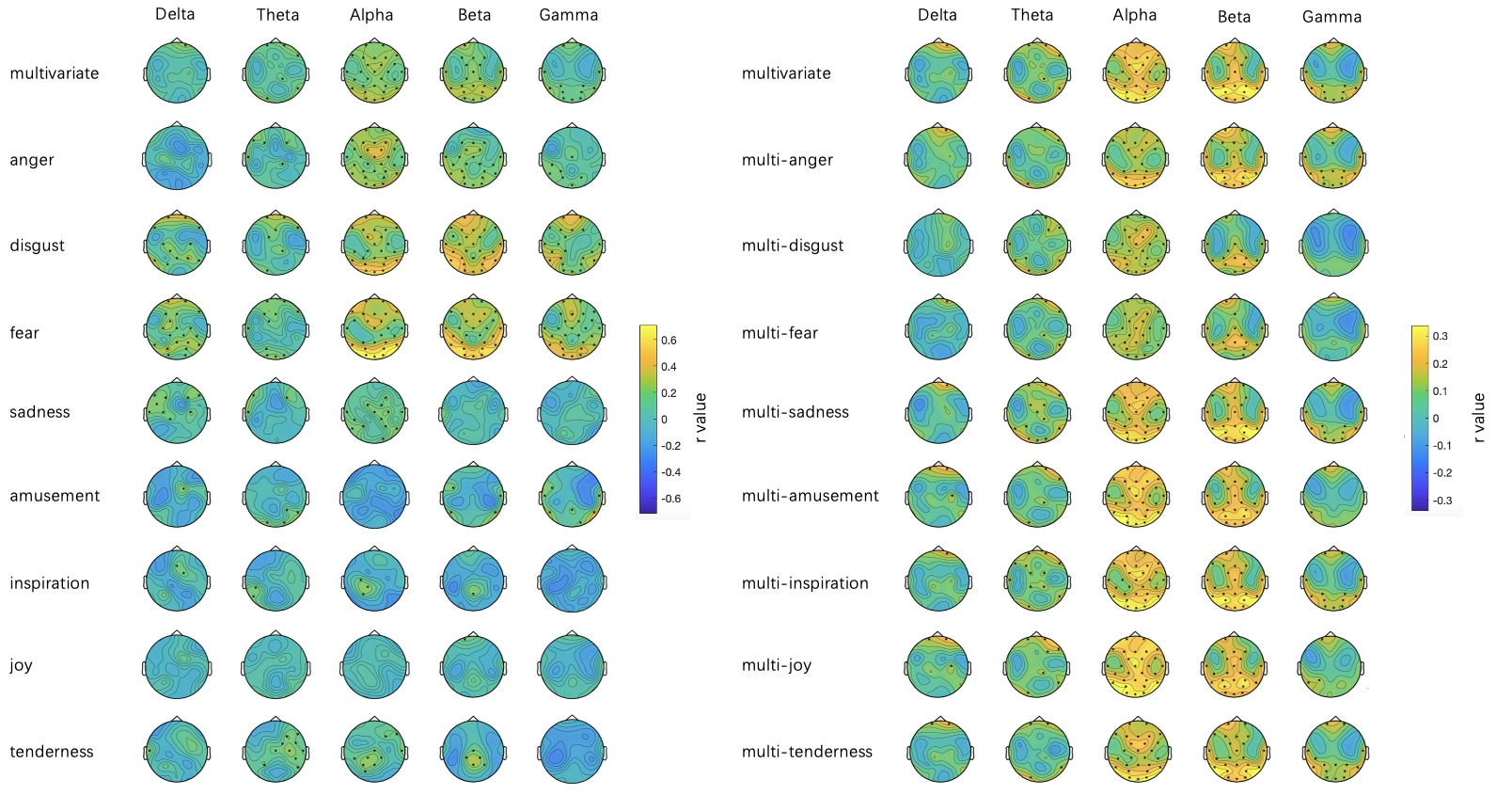

(a) (b)

**Figure S4.** (a) The inter-situation RSA correlation values between the multivariate /ablated multivariate emotion experience ratings (excluding each variate from the 8-variate respectively, e.g., multi-anger = the 7-variate without anger) and the PSD feature at each channel in the five frequency bands. (b) The inter-situation RSA correlation values between the multivariate /each univariate emotion experience ratings and the PSD feature at each channel in the five frequency bands. The channels with significant correlation values (ps < 0.05, FDR corrected) are marked in the topographies.

Note: The topographies corresponding to multivariate in the two subplots (the first row) are the same but with different colorbar scales.
